# Tandem repeats drive variation of intrinsically disordered regions in budding yeast

**DOI:** 10.1101/339663

**Authors:** Michael Babokhov, Bradley I. Reinfeld, Kevin Hackbarth, Yotam Bentov, Stephen M. Fuchs

## Abstract

Copy-number variation in tandem repeat coding regions is more prevalent in eukaryotic genomes than current literature suggests. We have reexamined the genomes of nearly 100 yeast strains looking to map regions of repeat variation. From this analysis we have identified that length variation is highly correlated to intrinsically disordered regions (IDRs). Furthermore, the majority of length variation is associated with tandem repeats. These repetitive regions are rich in homopolymeric amino acid sequences but nearly half of the variation comes from longer-repeating motifs. Comparisons of repeat copy number and sequence between strains of budding yeast as well as closely related fungi suggest selection for and conservation of IDR-related tandem repeats. In some instances, repeat variation has been demonstrated to mediate binding affinity, aggregation, and protein stability. With this analysis, we can identify proteins for which repeat variation may play conserved roles in modulating protein function.

## Introduction

Understanding how proteins carry out diverse functions and how these functions are regulated by the cell is a critical challenge of biology. Most proteins are produced as a linear polymer of amino acids that fold into three-dimensional structures and this structure is generally thought to determine protein function. Intrinsically disordered proteins (IDPs) or intrinsically disordered regions (IDRs) within proteins are sequences that generally do not adopt a single defined configuration. Recent evidence has now demonstrated that conformational disorder plays an important regulatory role in tuning protein interactions and stability (Habchi, Tompa et al., 2014, Tompa, 2012).

Our lab has been interested in a subset of IDRs that consist of repetitive amino acid sequences such as the repetitive C-terminal domain of RNA polymerase II Rpb1p (Babokhov, Mosaheb et al., 2018, Morrill, Exner et al., 2016). IDRs are generally enriched for amino acids that are not structure promoting (Campen, Williams et al., 2008). Curiously, these same amino acids are also often enriched in repetitive amino acid sequences in proteins (Simon & Hancock, 2009). A unique aspect of repetitive amino acid sequences is that they are often encoded by repetitive DNA. Repetitive DNA sequences are known to be genetically unstable, often resulting in expansions and contractions within the genomic sequence (Gemayel, Vinces et al., 2010, Richard & Dujon, 2006). Studies of repetitive regions have generally focused on trinucleotide repeat sequences (Albrecht & Mundlos, 2005, La Spada & Taylor, 2010) but our group recently showed that DNA encoding longer repetitive sequences also showed genetic instability (Morrill et al., 2016). The genetic diversity that could result from repeat instability is now being realized as a potential important player in complex traits. Additionally, previous studies have hinted at significant overlaps between repetitive sequences and IDRs (Jorda, Xue et al., 2010, Simon & Hancock, 2009, Tompa, 2003). Thus, the primary goal of the work described below was to determine whether repetitive sequences are a general feature of IDRs and how they might function to tune IDR function.

We hypothesize that genetic instability in repetitive regions generally contributes to population-level genetic variation. In this work, we analyzed existing genomic sequencing data from 93 *Saccharomyces cerevisiae* genomes (Strope, Skelly et al., 2015) and additional available data from related yeast species (Scannell, Zill et al., 2011) to determine whether repeat variation may be a significant contributor to the function of IDR domains. In brief, we found length polymorphisms in nearly 10% of yeast IDR domains. The vast majority of this variation derives from copy number variation in amino acid tandem repeats contained within these IDRs. Copy number variation is extensive and most commonly found in homopolymeric amino acid repeats and larger oligopeptide (>5 amino acid) repeat motifs. Lastly, variable repeats within IDRs are highly conserved across yeast species, suggesting an important biological function for these sequences. We propose that the genetic variation caused by repetitive sequences would further expand the regulatory features of IDRs in proteins.

## Results and Discussion

### A pipeline to identify IDR variation

Given the previously hinted association between tandem repeats and IDRs (Jorda et al., 2010, Simon & Hancock, 2009, Tompa, 2003), we developed a pipeline to characterize IDR and repeat variation in budding yeast (Figure 1). A list of IDRs was compiled using the VSL2B algorithm for 5,860 annotated genes to yield 7,531 predicted IDR sequences using a minimum IDR length of 30 amino acids to approximate long IDRs (Peng, Radivojac et al., 2006). High quality genomic sequences from the 100 yeast genomes project (Strope et al., 2015) were aligned and the predicted IDRs were scanned for length variation. IDRs were identified as either variable or non-variable and the presence of tandem repeats was noted and compared to predictions based on the XSTREAM algorithm (Newman & Cooper, 2007). The resulting dataset characterizes the variation and repetitiveness for all IDRs in our pipeline (supplementary Table S1 online). IDRs identified as both variable and repetitive were then further analyzed to determine selection and conservation.

**Figure 1.**
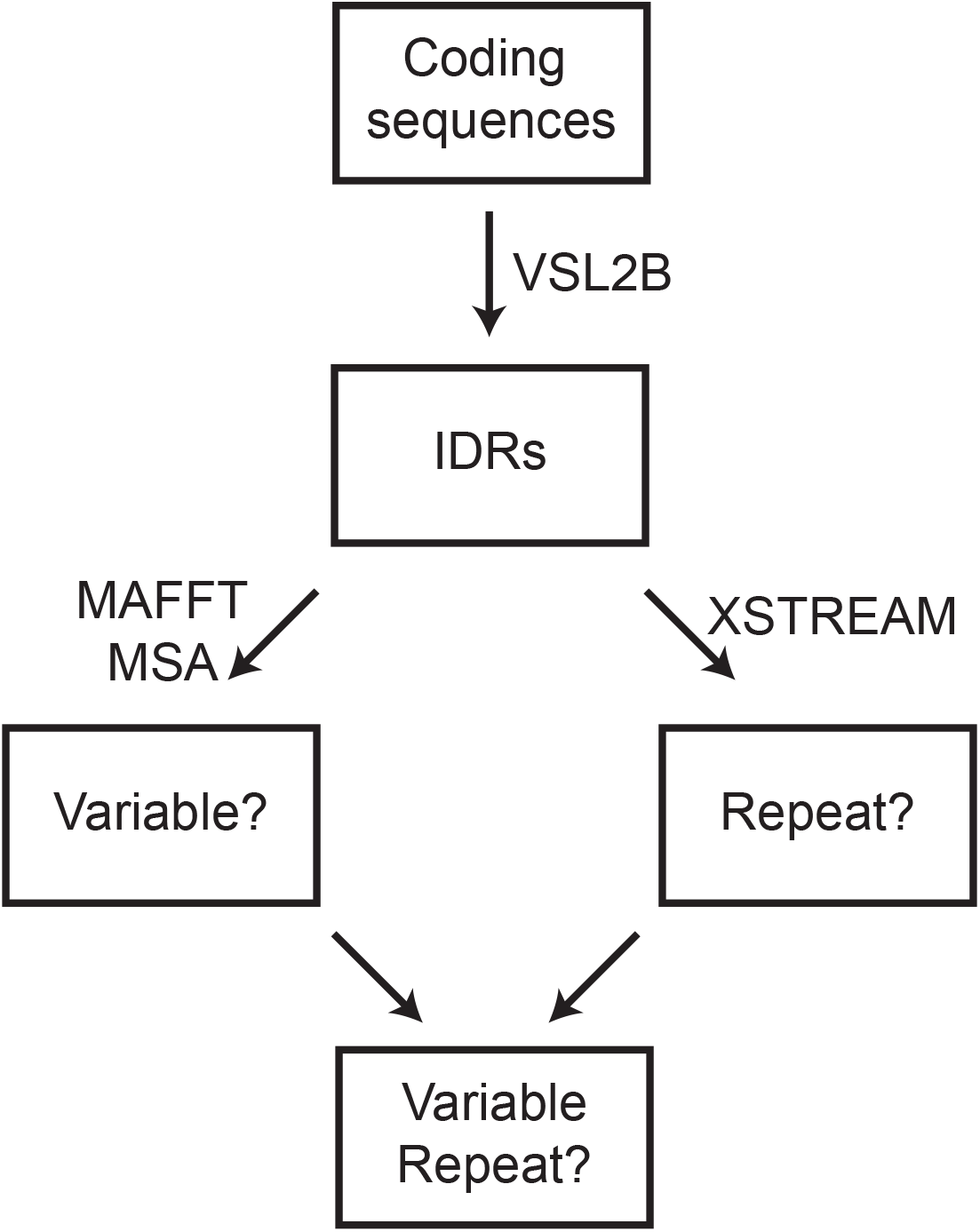
Flowchart of variable IDR identification. Coding sequences for all open reading frames of *S. cerevisiae* were inputted into VSL2B disorder predictor (Peng et al., 2006) to obtain a list of IDRs. The resulting sequences were aligned using MAFFT multiple sequence alignment (Katoh & Standley, 2013) to uncover length polymorphisms. In parallel, the repeat-finding algorithm XSTREAM (Newman & Cooper, 2007) was used to create a list of all repetitive sequences within IDRs. The overlap between these two data sets was then curated to identify variable repetitive sequences.

### IDR variation is associated with tandem repeats

Using the pipeline described above, we identified 913 IDRs that exhibited length polymorphisms in at least one of the 93 genomes examined (Figure 2A). Many of these polymorphic IDRs varied by only a single amino acid where others exhibited differences of more than 100 amino acids between the 93 examined sequences. Many proteins contained multiple IDRs and of these, 105 contained more than one polymorphic IDR. These findings are in line with previous studies that show genetic instability in IDR coding regions (Brown, Takayama et al., 2002, Nilsson, Grahn et al., 2011) and indicate the widespread variation that exists within natural and laboratory strains of budding yeast.

**Figure 2.**
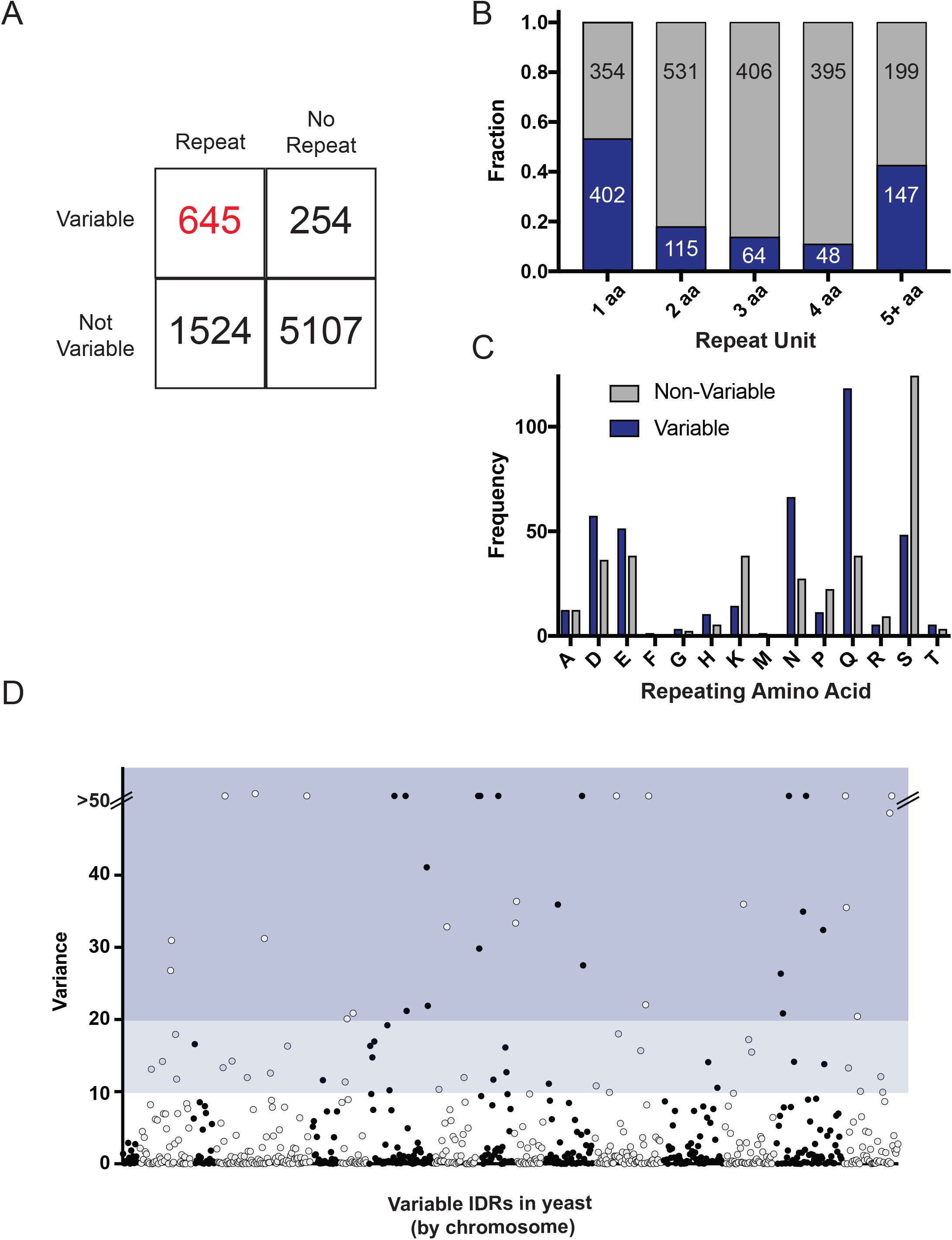
Characteristics of variable repeats in IDRs. **A**) Total numbers of identified IDRs based on repetitiveness and variability. **B**) Fraction of variable and non-variable repeats for each class of repeat unit length. **C**) Total numbers of variable and non-variable single amino acid repeats broken down by the repeated residue. **D**) Manhattan plot of length variances of variable IDRs by chromosome. The two shaded regions represent the top 5% (dark) and 10% (light) of IDRs with respect to variance.

IDRs and tandem repeats in proteins share many similarities including enrichment for small polar amino acids and high rates of genetic mutation. We therefore examined whether tandem repeat variation was responsible for the polymorphisms observed in IDRs. Using XSTREAM we identified 668 variable tandem repeats within the 913 polymorphic IDRs (~73%) (Figure 2A). This number is likely conservative as we identified tandem repeats as being at least five consecutive amino acids for homorepeats (Chavali, Chavali et al., 2017) and greater than two repetitions of larger repeats. Overlooked motifs with smaller copy numbers or sequence substitutions may represent degenerated repeats that could still retain functional significance. The remaining 240 variable IDRs were generally short duplications of sequence that were observed in a small subset of the 93 strains for a given IDR. This variation may represent genuine polymorphisms or may simply be an artifact of short-read next generation sequencing.

Close to half of all variable IDRs contained polymorphic homorepeats, commonly polyQ or polyN, which were highly polymorphic in length across the 93 strains. These homorepeats most frequently demonstrated poly-Q, -N, -D and -E sequences (Figure 2B and Table S1), in line with previous findings on both repeat and IDR-enriched amino acids (Campen et al., 2008, Gemayel et al., 2010). Repeat motifs of homorepeats were also differentially enriched between variable and non-variable IDRs. Homorepeats in variable IDRs were heavily skewed towards polyQ and polyN, while non-variable IDRs had many more polyS and polyK repeats (Figure 2C). Further analysis of these different enrichments would help uncover why particular repeats are variable within IDRs.

Trinucleotide repeats consisting of repeating sequences of a single codon are well known to be genetically unstable and we generally expected that this mechanism would be responsible for the majority of the homorepeat IDR variation genome-wide (Usdin, House et al., 2015). Interestingly, we uncovered several variable homorepeats that were not encoded by trinucleotide repeats signifying variation in these regions is governed by more complex mechanisms than was previously suggested. Table S2 shows the codon distribution for long and short forms of polyQ repeats (Table S2). In addition to these low-complexity homorepeats we found that repeating sequences made up of more than a single amino acid comprised nearly half of the variable repeats (Figure 2B). Larger motifs containing five or more amino acids were most frequently found to be variable in length, while motifs of two to four amino acids were more likely to be non-variable. Finally, we also identified small linear duplications that created new repeating motifs leading to IDR length variation (Table S3). These sequences may be examples of nascent repeats and it would be interesting to further explore these motifs to determine whether there are specific genomic features that contribute to the genesis of new repeats.

As stated above, we found many IDRs for which there was a wide variance in length, due to polymorphism in a repetitive region. In some cases we also found several length polymorphisms but they were restricted to just a few individuals. We therefore calculated the sample variance of repeat length to represent both the repeat length polymorphisms and the frequency of a given variant within the population (Table S1). Visualizing the data on a Manhattan plot shows the variances for all 668 variable tandem repeat IDRs across the 16 chromosomes of budding yeast (Figure 2D). Any variation in an IDR may be relevant for protein function but we stratified this data to highlight the IDRs with the greatest variance (top 5 and 10% as shaded blue boxes in Figure 2D). To further ensure the variation reported in the next-generation sequences were true representations of the genomic sequence we PCR amplified repeat regions and subjected them to Sanger sequencing. As an illustrative example, we present the variation of the C-terminal repeat of the *MNN4* gene (Figure 3). *MNN4* is known to be polymorphic in populations (Carvalho-Netto, Carazzolle et al., 2013) as seen by PCR amplification of genomic DNA, and encodes a protein important for mannosylphosphorylation of yeast cell wall proteins (Kim, Kang et al., 2017) and represents an interesting class of polyampholyte IDR domains (Das & Pappu, 2013, Sickmeier, Hamilton et al., 2007). Alignment of the resulting Sanger sequencing revealed extensive rearrangements of the repetitive motif and residue substitutions among four sample stains (Figure 3B), with the overall distribution of copy number variation seen in Figure 3C. We have performed similar analysis on several other IDRs, confirming to us that the variation reported in the next-generation sequence data largely represents true genetic polymorphism and not errors from sequencing or genome assembly.

**Figure 3.**
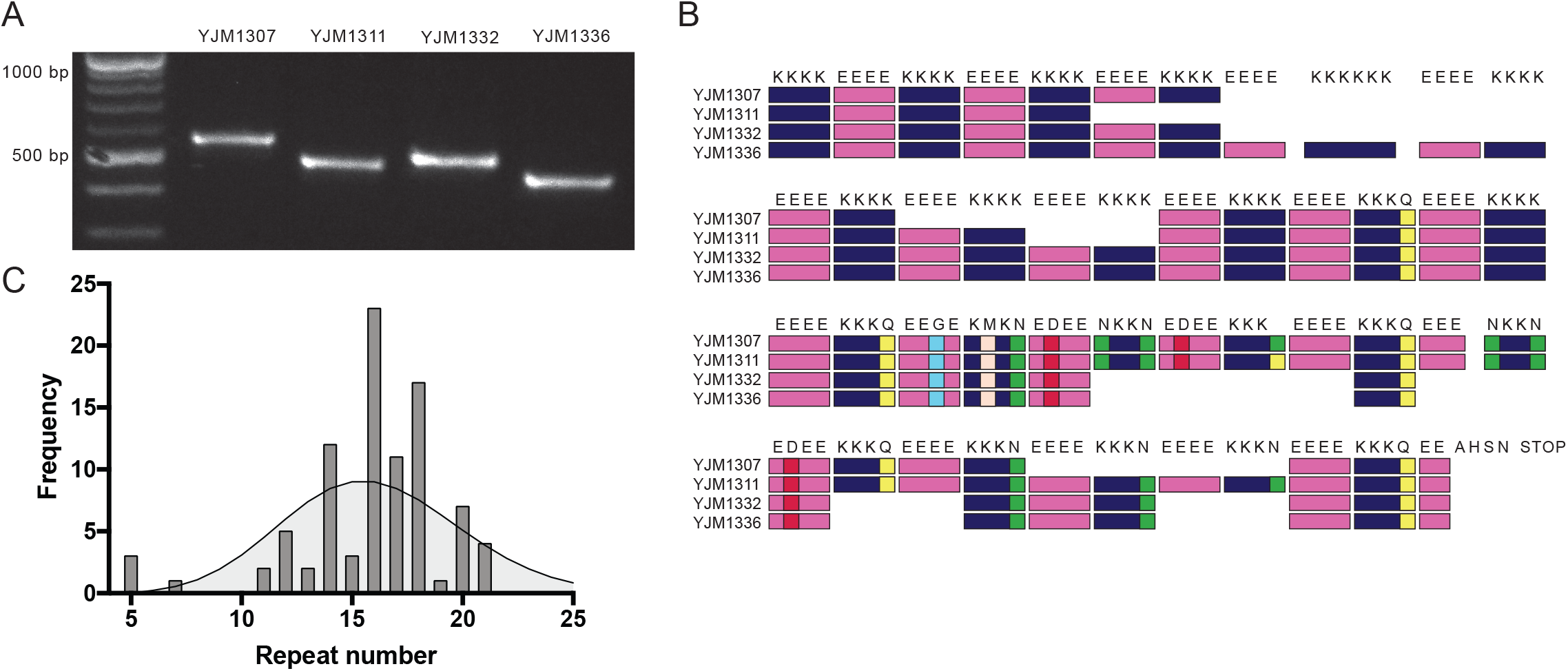
Variation in the C-terminal repeat of *MMN4*. **A**) Agarose gel of an amplified region of *MMN4* showing the length variation in the C-terminal repeat region of four representative strains. **B**) Diagram of *MMN4* repeat structure alignments derived from Sanger sequencing of the PCR products from A. **C**) Frequency distribution of all 93 *MMN4* repeats organized by repeat copy number. The curve represents a theoretical Poisson distribution calculated from the observed mean copy number.

### Variable and repetitive IDRs are conserved across Saccharomyces species

We and others have proposed that variation in repetitive regions is an important driver of protein diversification (Fuchs, 2013, Gemayel, Yang et al., 2017, Morrill et al., 2016, Rogers, McConnell et al., 2017, Verstrepen, Jansen et al., 2005, Zhao, Strope et al., 2014). However, it is also possible that some or many of these repeats have randomly risen within the genome and thus are unlikely to contribute to protein function. Conservation of a repeat across different species would suggest it plays some functional role. To test this assertion, we examined evolutionary conservation of variable repeats across four closely related species of the *Saccharomyces sensu stricto (SSS)* genus using existing high quality genome data (Scannell et al., 2011). We found that 573 of 668 variable IDRs (~85%) were found in at least one other species of the SSS. Such a high level of conservation argues that the repeat regions within IDRs are not the consequence of random mutagenesis but have important biological functions within budding yeasts. In addition to the conservation of the repeat sequence, we also observed copy number differences of tandem repeats across the SSS, indicating that repeat variation may play an important role in protein function across species as well as evolution/speciation (Table S3). However, we currently don’t have enough representative samples to know the full range of repeat lengths that might exist for a given IDR in these other species. Further genomic sequencing and long-read next generation sequencing of *sensu stricto* strains and other fungi will be required to fully appreciate tandem repeat variation across evolutionary time.

In addition to completely conserved repeats, we found 18 IDRs that preserved the tandem repeat nature of the loci while having a different consensus sequence. A prime example of these repeats is located in an IDR of Pan1, where the 6-mer PIQPVQ repeat in *S. cerevisiae* is replaced with either a PAQ or PVQ trimer motif in other *sensu stricto* organisms (Supplementary Figure 1D). In all 18 cases, the difference in repeat motif between *SSS* sequences are conservative and thus we predict them to not differ greatly in function. We also identified 31 repeats that did not have corresponding sequences in the other *sensu stricto* genomes, and thus were novel to *S. cerevisiae*. All but three of these non-conserved repeats contained repeating motifs with longer periods (2 or more amino acids) and were absent from the other *SSS* species (Figure S1). These nonconserved repeats may have been acquired recently or may be required for an adaptation specific to *S. cerevisiae*. Lastly, we note that we were unable to characterize 52 variable repeats because the corresponding genes were absent in the *SSS* genomic sequences (Table S4). Overall, our analysis of tandem repeats across the *SSS* demonstrates that variable repeats located within IDRs are highly conserved in budding yeasts, implicating them in biologically relevant functions.

### Potential role for genetic selection in repetitive IDRs

Tandem repeat variation is thought to be a mechanism for rapid evolution of protein coding sequences (Marcotte, Pellegrini et al., 1999). As many variable IDRs are conserved, this suggests they may play some role in protein function. Therefore, we might expect two scenarios: the first is a situation where there is genetic selection for repeat length such that a few discrete repeat lengths would dominate the population. This might be the expectation of an IDR that functions to link two folded domains, for example, where the length is highly constrained by the function of the neighboring domains. The second scenario would be one where length polymorphism is neutral or even evolutionarily advantageous. In this scenario, we would expect to see some distribution of repeat lengths for the sample tested. We looked at the distribution of repeat lengths for each gene and determined that nearly all genes conformed to the first scenario, *i.e*. they showed variation but a high level of genetic selection for a preferred repeat length. To illustrate this, we present the length distribution of two orthologs in yeast which both harbor variable-length repetitive IDRs in their C-terminal domain, *VHS3* and *HAL3* (Figure 4). *VHS3* is among the most highly polymorphic IDRs we examined with repeat lengths ranging from 21 to 64 consecutive aspartic acid residues (Figure 4A) whereas *HAL3* had, on average, a much shorter polyD region and the distribution of lengths was more tightly clustered (Figure 4B).

**Figure 4.**
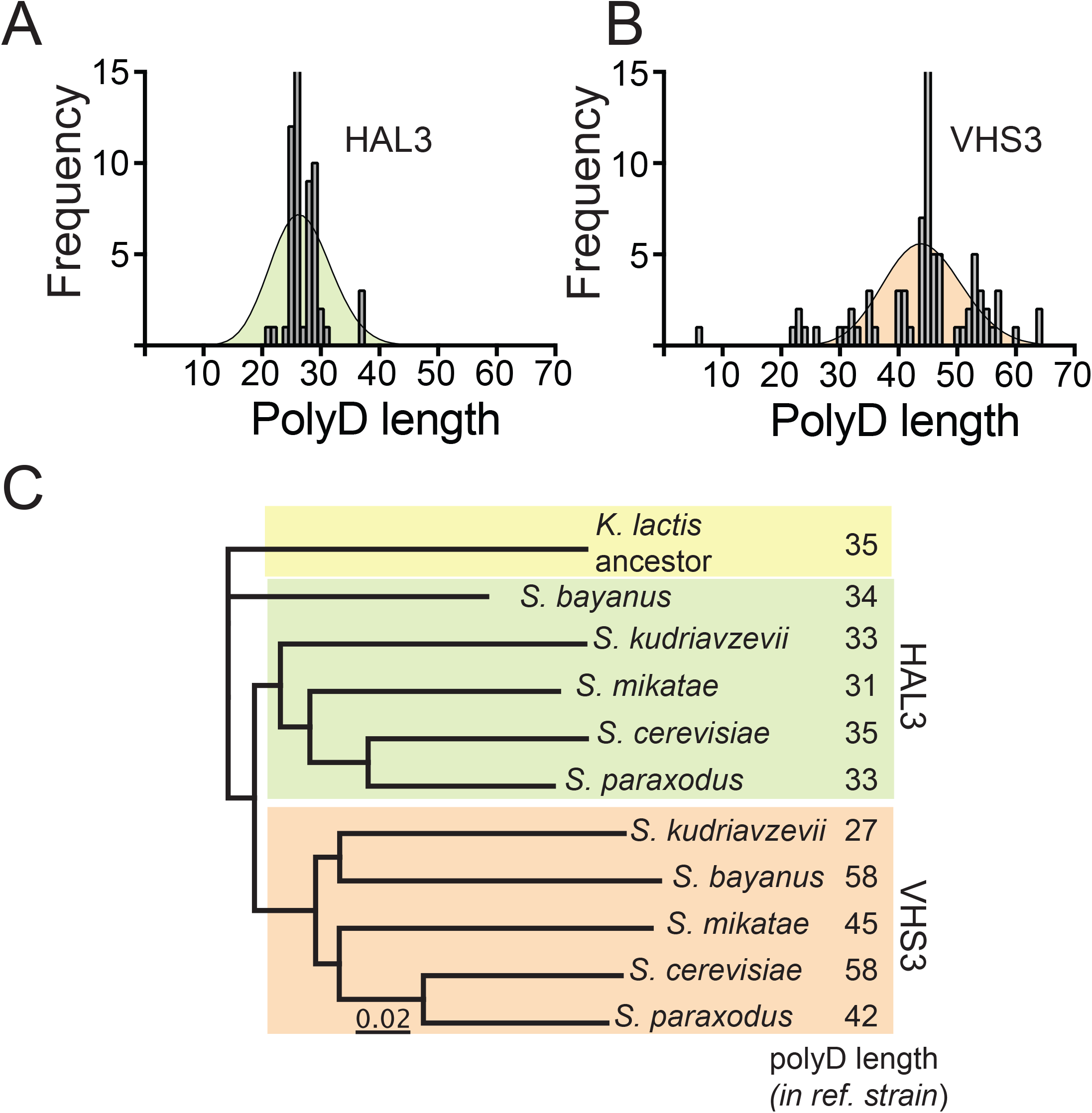
Comparison of polyD variation in paralogs *HAL3* and *VHS3*. **A**) Frequency distribution of HAL3 polyD homorepeat for all 93 yeast strains together with the theoretical Poisson distribution curve. **B**) Same frequency distribution for the VHS3 polyD homorepeat. **C**) Phylogenic tree of genetic distances between the *Saccharomyces sensu stricto* species and the pre-whole genome duplication *Kluyveromyces lactis* ancestor. Distances were calculated based off of changes to *HAL3* and *VHS3* polyD homorepeat copy number listed to the right.

In order to estimate selective pressures on these two IDRs we compiled the polyD copy number for all of the *sensu stricto* yeasts and used repeat variation as a proxy for genetic distance between the species and *Kluyveromyces lactis*, an evolutionary ancestor that precedes the whole genome duplication seen in the *sensu stricto* clade (Wolfe & Shields, 1997). We saw that the polyD repeat encoded by both *VHS3* and *HAL3* is found in all organisms, including *K. lactis*. Overall, we found that the *VHS3* and *HAL3* variation among the SSS species recapitulated the pattern in *S. cerevisiae*: the *HAL3* polyD was tightly clustered while the *VHS3* polyD showed greater variation (Figure 4C). Interestingly, the genetic distances that we established based on repeat variation closely resembled the overall phylogenic relationship between the *sensu stricto* (Scannell et al., 2011), suggesting that variation in IDR repetitive sequences tracks with evolutionary changes.

## Conclusion

In summary, we have uncovered extensive IDR length polymorphisms across 93 wild and laboratory strains of *S. cerevisiae*. The majority of the length differences between strains can be explained by copy number variation in both single amino acid and oligopeptide tandem repeat sequences. We have exhaustively characterized the variation that we observed and presented several illustrative examples of the trends within repetitive variable IDRs. Two key patterns have emerged as a result of our analysis. The first is that repeat variation is more complex than was previously appreciated, covering many different kinds of motifs at both the DNA and the protein level. The second is that although repetitive sequences are variable within *S. cerevisiae*, they are highly conserved across budding yeast species, arguing for an important biological function for the repeats and perhaps their copy number variants. Consequently, our analysis of repeat length variation provides a foundation to further study both the mechanisms that lead to variation within IDRs and investigate the impact of length variation on protein function.

## Methods

### Prediction of disordered and tandem repeat regions

Variation within intrinsic disordered regions was measured using the data available from 93 recently sequenced *S. cerevisiae* genomes (Strope et al., 2015). Genomic sequences were acquired and the open reading frames of 5,860 annotated genes were aligned in Geneious v. 10.2.3 [Biomatters Ltd.] using MAFFT v. 7.308 with the default settings (Katoh, Misawa et al., 2002, Katoh & Standley, 2013) and corrected manually when necessary. From this data set we removed regions associated with retrotransposons and dubious open reading frames. Disordered regions in proteins were compiled using the VSL2B disorder prediction algorithm using *S. cerevisiae* reference genome release 63.3 Saccharomyces Genome Database and was restricted to regions that were at least 30 amino acids in length (Oates, Romero et al., 2013, Peng et al., 2006). Each IDR was given a unique identifier based on its start position and assigned as either variable or nonvariable based on visual inspection of the alignments.

Tandem repeats were acquired using the XSTREAM repeat prediction algorithm (Newman & Cooper, 2007). We used XSTREAM parameters were set as: minimum character identity, *I* = 0.7; minimum consensus match, *I* = 0.8; maximum consecutive gaps, *g* = 3; minimum period, MinP = 1; minimum length, L = 5; any other settings were set to default. These settings were chosen to be very inclusive in order to identify both short perfect repeats and longer degenerate repeats in the reference proteome. A minimum overall length for the repeat region was set at five amino acids (Chavali et al., 2017). Using the data from XSTREAM each IDR was assigned as either containing a tandem repeat (Y) or not (N). Note in supplemental tables that some longer IDRs contained more than one tandem repeat region as called by XSTREAM. Variation in each unique repeat was recorded separately (Table S1).

### Assessment of IDR variation

The presence or absence of repeat length variation at each individual IDR was assessed using the MAFFT alignments of the 100 yeast genomes. Alignments of each gene in the Geneious browser were examined manually for the presence of gaps that indicated sequence variation between the different yeast strains. IDRs that had at least one gap within their sequence range were scored as variable and the number of variable regions and the variable protein motifs were recorded. Only gaps in the alignment from insertions and deletions (indels) were examined. Variation due to single nucleotide substitution was excluded from the current study. The range of variation present at a given IDR was calculated as the difference in amino acid length between the longest and shortest forms of the variable sequence. We also calculated the length variance for each variable IDR. Consequently, the repetitiveness of each variable IDR was also annotated as either repetitive or not and the number repeats per IDR as well as their sequences were noted. A number of highly variable repeats were realigned using nine kilobases of flanking sequence in order to get accurate frequency counts. Even after realignment, a subset of 31 hypervariable repeats still did not yield sufficient alignments to get frequency data and were marked as variable but excluded from further analysis (Table S5). The frequency data for each variable repeat was plotted and compared to a theoretical Poisson distribution for an equivalently sized dataset with mean in GraphPad prism by a chi-squared test. Variable tandem repeats were then categorized as either having a Poisson or a non-Poisson distribution.

### Determination of variable repeat conservation across Saccharomyces sensu stricto

Alignments of annotated genes from high-quality genomes of four Saccharomyces species closely related to budding yeast (Scannell et al., 2011) were downloaded and compared to the alignments from the 100 yeast genomes resource. The corresponding positions of each IDR identified as variable in *S. cerevisiae* were examined manually in the alignments of the other four species to look for variable repeats. If the tandem repeat was present in both the *S. cerevisiae* and at least one of the *sensu stricto* alignments, then the repeat was classified as conserved. Occasionally, the repeat sequence was not conserved but there was a different sequence that was still repetitive at the corresponding location. In these cases, the repetitiveness was classified as conserved while the sequence was not. A subset of variable repeats could not be classified because there was no corresponding alignment file in the *sensu stricto* dataset.

## Acknowledgements

The authors would like to thank J. Michael Reed for many discussions on statistical methods and all the members of the Fuchs lab for helpful discussions and assistance in analyzing repeat variants. This work was funded in part by Army Research Office Grant W911NF-16-1-0175 and a Tufts Collaborates grant to S.M.F.

## Author contributions

M.B., B.I.R., and S.M.F. were responsible for the initial design and execution of the analyses. All authors were responsible for analyzing the data. M.B. and S.M.F. wrote the manuscript and prepared the figures. All authors helped edit the manuscript.

## Supplemental Figure Legends

**Figure S1.**
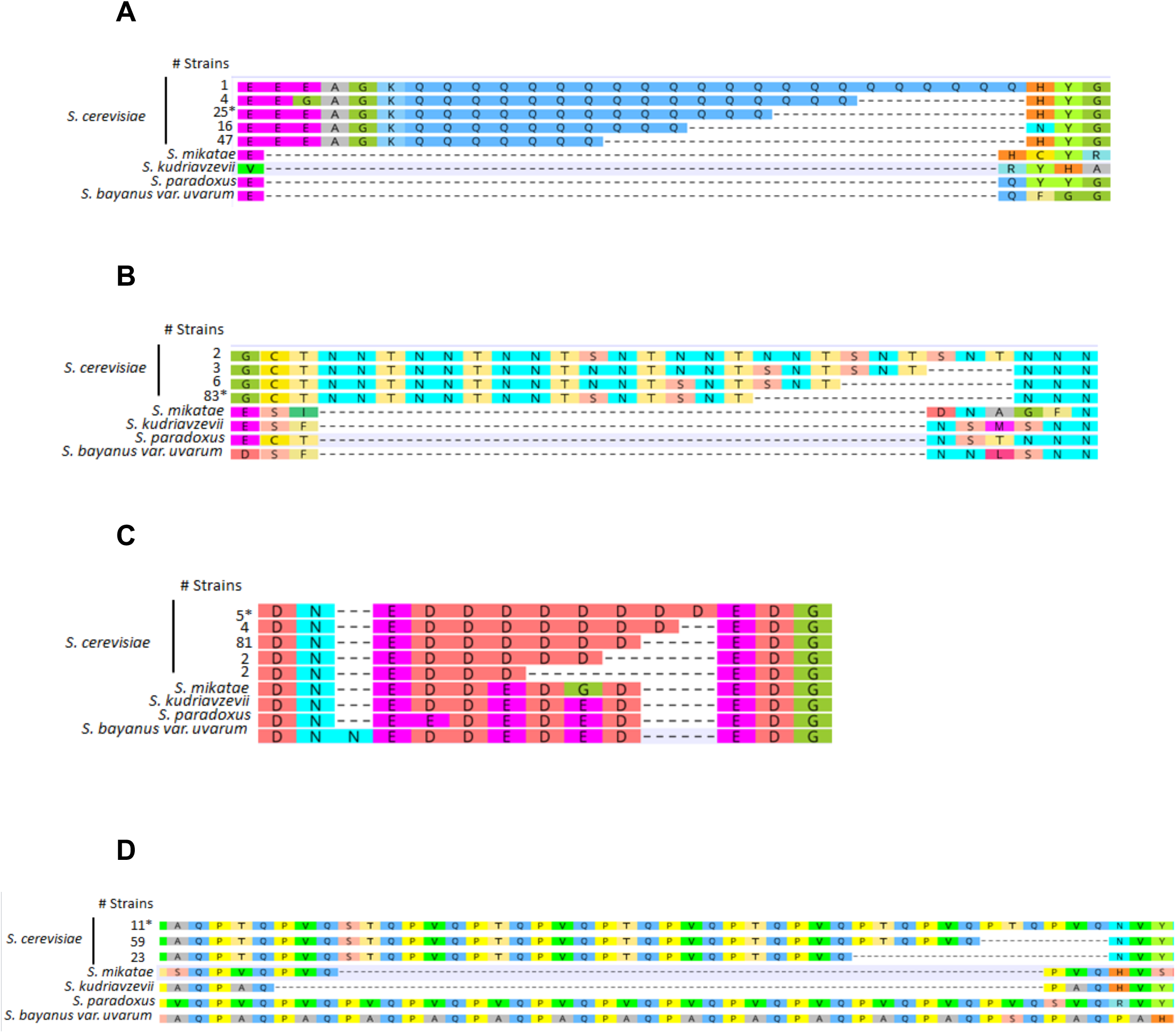
Examples of non-conserved and sequence-variable repeats in the *Saccharomyces sensu stricto*. **A**) MAFFT alignment of the five length variants of the JJJ2p polyQ homorepeat in *S. cerevisiae* with the corresponding sequences in the SSS that lack the repeat. The number of *S. cerevisiae* strains for each copy number variant is listed to the left with the starred number representing the copy number in the S288C reference genome. **B**) Alignment of the *YPR003C* NNT repeat. **C**) Alignment of the *KAP104* polyD homorepeat in *S. cerevisiae* showing a mixed polyD/polyE repeat in the other SSS species. **D**) Alignment of the *PAN1* repeat showing different consensus sequences of the repeat depending on the species of the *Saccharomyce sensu stricto* that is examined.

